# Effects of Inflammatory Factors on the Migration and Repair of Intestinal Mucosal Cells at Pancreaticojejunostomy Anastomosis: An Experimental Study in Beagle Dogs

**DOI:** 10.1101/2025.02.26.640054

**Authors:** Shixing Wu, A Tigu, Bolin Zhang, Cheng Wang, Jiangang Li, Junxiang Zhang, Shouchao Zhang, Cheng Geng, Xinjian Xu

## Abstract

**Objective:** To investigate the effects of inflammatory factors on the migration and repair of intestinal mucosal cells at the site of pancreaticojejunostomy (PJ) anastomosis and to observe the safety and feasibility of PJ with mucosal-priority healing (PM-PJ).

**Materials and methods:** Thirty-six Beagle dogs were randomly divided into two groups: the PM-PJ group (n=18) and the control group (n=18), which included classic end-to-side invagination pancreaticojejunostomy (IPJ). Six Beagles were sacrificed on the 7th, 14th, and 21st days after surgery to obtain pancreaticojejunostomy anastomosis tissues. The primary outcomes were postoperative mortality, morbidity, and pathological changes in the anastomosis. The secondary outcomes were the expression levels of the cytokines TGF-β1, IL-10, TNF-α, IL-6, collagen, and α-SMA.

**Results:** All surgeries were successfully performed. In the study group, the incidences of anastomotic leakage and pancreatitis were 0% and 11.1% (2/18), respectively, whereas in the control group, the incidences were 5.6% (1/18) and 16.7% (3/18), respectively. The levels of serum TGF-β1, IL-10, TNF-α, and IL-6 were increased at all time points after surgery in both groups, and these increases were more significant at certain time points in the study group. Pathological observation revealed that the degree of anastomotic healing in the study group was greater than that in the control group at all time points. The protein expression levels of collagen I/III and α-SMA in the anastomoses of both groups changed over time, and the changes were more significant at certain time points in the study group.

**Conclusion:** Inflammatory factors play a positive role in the early stage of PJ anastomosis healing by stimulating collagen secretion and synthesis. PM-PJ is a safe and simple surgery with good anastomotic healing and a low incidence of complications.

## 1. Introduction

In the past few decades, substantial clinical progress has been made in pancreatoduodenectomy (PD), which has provided patients with the best long-term survival opportunities. However, even in high-volume centres, 40%-60% of patients experience postoperative complications and 2%-5% of patients are at risk of in-hospital death [1]. The difficulty of PJ, a key step in PD, lies in the soft texture and rich blood supply of the pancreas. Serious complications such as pancreatic fistula, bleeding, and death are likely to occur after surgery [2], and thus its safety remains a challenge and is directly related to the success of the surgery as well as the patient’s postoperative recovery. A key research area of PJ is the necessity of duct-to-mucosa anastomosis [3], which can ensure the direct flow of the main pancreatic duct into the intestine; this maintains the continuity of the pancreatic duct and jejunal mucosa and facilitates tissue healing, which is consistent with the normal physiological process.

During the healing process of PJ anastomosis, repair of the intestinal mucosa mainly depends on the continuous proliferation and differentiation of cells in the crypts of the intestinal glands and the migration of basal cells to form new basal cells [4]. At the same time, the migration and repair of intestinal mucosal cells cannot be considered independently of the extracellular matrix (ECM) [5]. Collagen is an important molecule in the ECM, as it not only provides mechanical support for tissues but also participates in various physiological and biochemical behaviours such as cell proliferation, migration, and signal transduction [6]. Multiple cytokines and signalling pathways, such as interleukin-6 (IL-6) [7], transforming growth factor-β (TGF-β) [8], and platelet-derived growth factor (PDGF) [9], play multiple roles in different stages of wound healing.

In clinical practice, through analysis of pathological sections of the anastomosis during re-pancreaticojejunostomy in patients with pancreatic duct stenosis after PD, our team reported that the healing process of the PJ anastomosis can be divided into two connections: a mucosal connection (inner layer) and a mechanical connection (outer layer). The inner layer involves healing of jejunal mucosal epithelial cells and pancreatic duct epithelial cells, while the outer layer involves fibrous tissue healing. Regardless of the PJ method, the final connection is between the intestinal mucosa and the epithelial cells of the pancreatic duct. Different anastomosis methods result in different crawling distances of the intestinal mucosa. Starting from the crawling distance of the intestinal mucosa, we propose the theory of “mucosal-priority healing during the PJ anastomosis healing process”. On the basis of this theory, our research team has performed a series of PJ surgeries. The purpose of this study was to evaluate the role of inflammatory factors in the healing of PJ anastomosis through histological analysis of PJ in Beagles.

## 2. Materials and methods

### 2.1. Establishment of the Experimental Animal Model

Healthy 12-month-old female Beagle dogs with a body weight of 9 ± 1 kg were used. Before surgery, the dogs were fasted for 6 hours and deprived of water for 12 hours. Before surgery, each dog was intravenously injected with 250 mg of cefotaxime sodium as an antibiotic prophylaxis. After induction with an intravenous injection of ketamine (2 mg/kg), tracheal intubation and assisted ventilation were performed with a ventilator. Enflurane was inhaled, and propofol was intravenously injected at a rate of 5 mg/kg/h to maintain anaesthesia.

Venous access was established through the right jugular vein, and vital signs were monitored during the surgery. Throughout the experiment, the animals were fed according to their body weight by dedicated personnel at the animal experiment centre, and water was provided ad libitum. The dogs were each housed in a single cage in a well-ventilated indoor environment with the room temperature controlled at 18-25°C and a relative humidity of 40%-60%.

A midline laparotomy incision was made in the upper abdomen. The distal jejunum was transected 15–20 cm distal to the ligament of Treitz and sutured intermittently with absorbable sutures. The proximal jejunum was anastomosed end-to-side with the distal end of the transected jejunum to ensure the patency of the gastrointestinal tract. The main pancreatic duct was carefully dissected and transected at the junction of the descending part of the duodenum and the pancreatic duct. The proximal end was ligated, and the other end was dissected and wrapped with pancreatic duct tissue for the PJ anastomosis. A pancreatic duct stent was placed in the intestine. After PJ, the surgical area was checked for bleeding, after which the abdominal incision was sutured.

### 2.2. Experimental Grouping and Surgical Methods

For the animal experiments, 36 dogs were randomly divided into two groups: the study group (n=18) underwent pancreaticojejunostomy with mucosal-priority healing (PM-PJ), and the control group (n=18) underwent classic end-to-side invagination pancreaticojejunostomy (IPJ). The pancreas in dogs is a V-shaped organ (Figure 1) consisting of the left lobe, right lobe, and pancreatic body. The diameter of the pancreatic duct is usually small at approximately 1–3 mm. PJ in 9-kg Beagles is technically difficult because the diameter of the transected pancreatic duct is only approximately 2 mm. In the study group, the jejunum was intermittently sutured to 2 stitches at the upper and lower edges of the pancreatic stump to ensure that the jejunum and the pancreatic stump were in close proximity. Then, an opening similar to the one in the pancreatic duct was made in the jejunum, and the seromuscular layer of the jejunal opening was sutured to the pancreatic stump centred on the pancreatic duct. The anterior and posterior walls of the anastomosis were sutured with 5/0 or 4/0 silk threads using 3–5 stitches, respectively. A stent was then placed in the pancreatic duct. It was not necessary to achieve perfect mucosa-to-mucosa anastomosis between the pancreatic duct and the jejunum. For small pancreatic ducts, to avoid tearing, it was only necessary to facilitate close migration and connection between the intestinal mucosa and the pancreatic duct epithelium (Figure 2). In the control group, when performing PJ, an appropriate position was selected to incise the full thickness of the jejunum. The size of the incision was the same as or slightly smaller than the diameter of the pancreatic stump. The pancreatic stump that had been inserted into the jejunum was sutured intermittently in a single layer. The posterior wall of the pancreatic capsule and the seromuscular layer of the jejunum were sutured intermittently with silk threads. The pancreatic stump was subsequently anastomosed with the full thickness of the jejunum in a circle. Finally, the anterior wall of the pancreatic capsule and the seromuscular layer of the jejunum were sutured intermittently. At a distance of 0.5–1 cm from the anastomosis, the seromuscular layer of the jejunum and the pancreatic parenchyma were continuously buried and sutured. On the 7th, 14th, and 21st days after surgery, 6 dogs in each group were sacrificed to obtain PJ anastomosis tissues for corresponding laboratory tests. The primary outcome index was anastomotic healing after surgery. The secondary outcome measures were postoperative mortality and morbidity.

**Figure 1:**
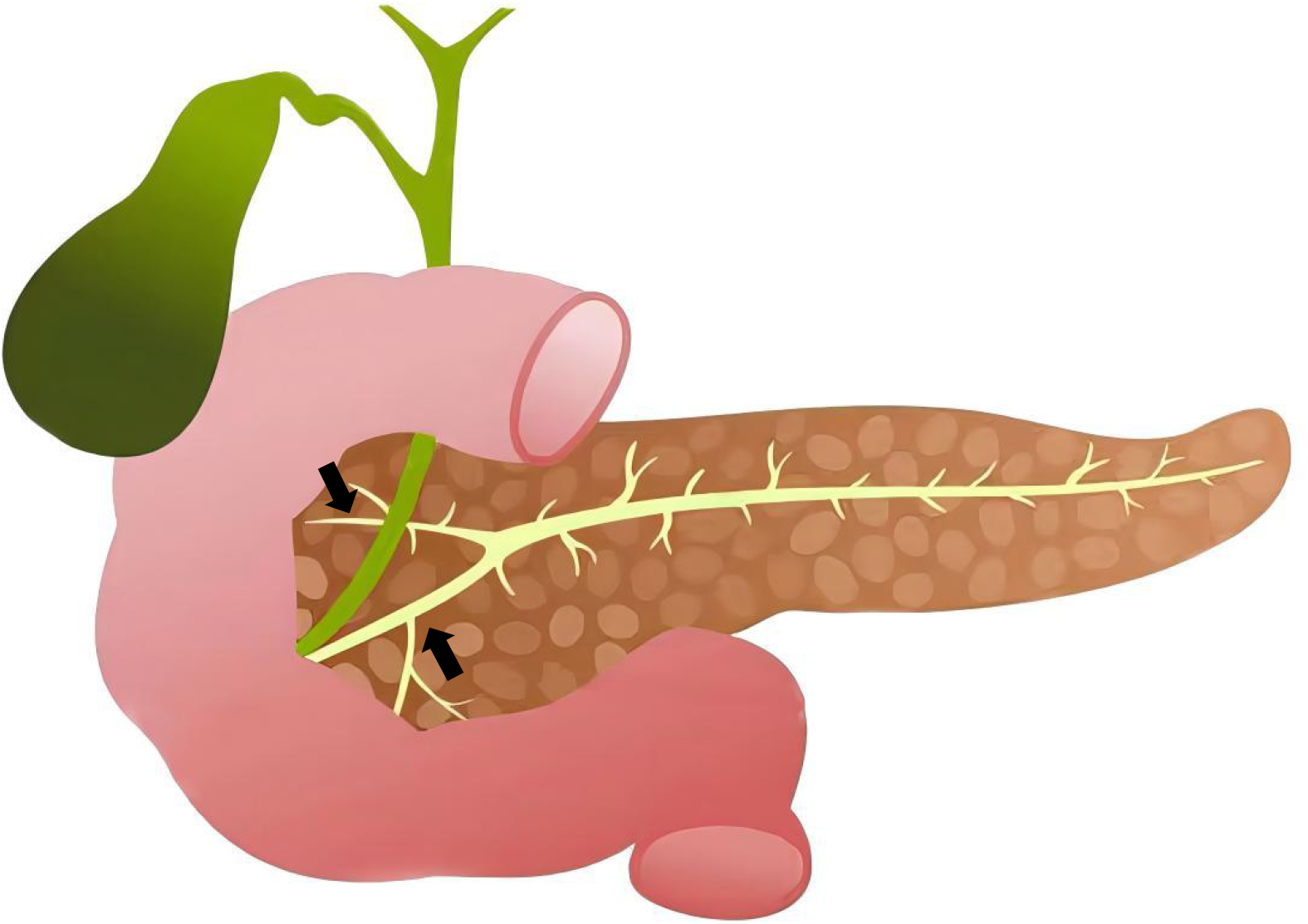
Anatomy of the pancreas in Beagles. The pancreas is divided into three parts: the left lobe, the right lobe, and the pancreatic body. Arrow heads: Position of the pancreatic duct.

**Figure 2.**
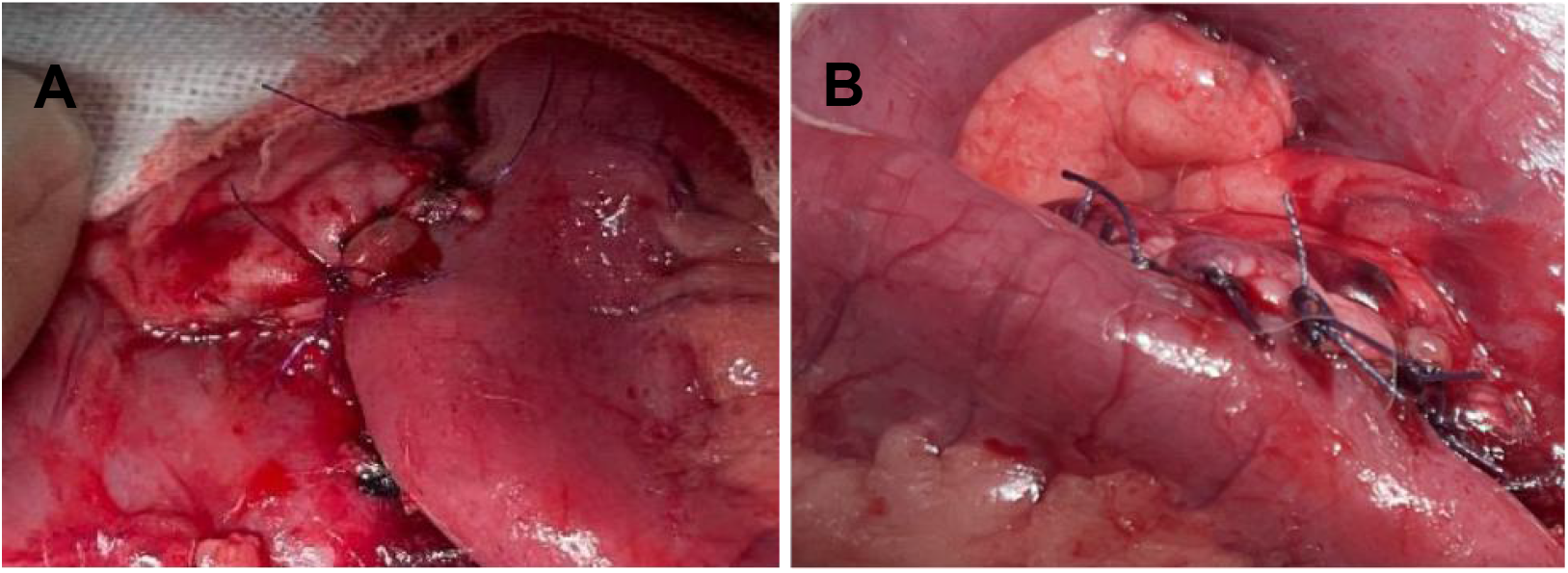
Beagle Pancreatic Intestinal Anastomosis Model: A, Study group (PM-PJ); B, Control group (IPJ).

This study was approved by the Ethics Committee of Xinjiang Medical University (Approval No. IACUC-20210303-55). All animals received humane care in the experiments conducted at the animal laboratory of Xinjiang Medical University in accordance with the Declaration of Helsinki for Animal Science Experiments (1975).

### 2.3. Observation Parameters

Peripheral venous blood was drawn before surgery and before sacrifice to measure fasting blood glucose and serum amylase levels. During the surgical procedure, the texture, shape, size of the pancreas, diameter of the pancreatic duct, and anastomosis time were recorded. The general conditions of the experimental animals after surgery were recorded, and the healing of the anastomosis at each postoperative stage was observed by gross pathology. The anastomosis tissues at each postoperative stage were fixed in 4% neutral paraformaldehyde for 24 hours and embedded in paraffin. The paraffin-embedded samples of the anastomosis tissues were cut into 4-mm sections for H&E staining, immunohistochemistry, and immunofluorescence staining.

The concentrations of TGF-β1, IL-10, TNF-α, and IL-6 in the serum were detected via enzyme-linked immunosorbent assay (ELISA). The serum samples obtained from the dogs were stored at room temperature for 2 hours or at 4°C overnight. To measure the contents of TGF-β1, IL-10, TNF-α, and IL-6 in the serum, 100 μl of serum or standards of different concentrations were added to each well of the microplate. After dilution, the microplate was covered with sealing film and incubated for 2 hours. After washing, enzyme-linked polyclonal antibodies specific for TGF-β1, IL-10, TNF-α, and IL-6 were added to the wells in the microplate, which was subsequently incubated at room temperature for 2 hours. After the samples were washed again with washing buffer, the colour intensity was measured at 450 nm in a microplate reader.

### 2.4 Data Analysis

The measurement data of the observed outcome indicators are presented as the means ± standard deviations. Count data are presented as percentages. Appropriate t tests, analyses of variance, or tests were used. P<0.05 was considered statistically significant. The data were analysed using Graphpad Pism9.5 statistical software.

## 3. Results

### 3.1. General Findings in the Experimental Animals

No significant differences were observed in the average body weight, fasting blood glucose, or serum amylase levels between the dogs in the two groups before surgery. No significant differences were observed in the average pancreatic duct diameter, pancreatic texture, or PJ anastomosis time between the two groups (Table 1). The serum amylase levels of the dogs in both groups increased within 7 days after surgery. From 7 to 14 days after surgery, the levels in both groups tended to decrease. From 14 to 21 days after surgery, the levels in the study group showed a slow upward trend, whereas the trend in the control group was flat, with no significant change. The serum amylase levels of the two groups were the same but were higher than the preoperative values (Figure 3).

**Table 1.** Comparison of general surgical data between the two experimental groups of Beagle dogs.

|  | Study group (n=18) | Control group (n=18) | *p* |
| --- | --- | --- | --- |
| Weight (kg), mean (± SD) | 9.08 ± 0.56 | 9.03 ± 0.36 | 0.787 |
| FBG (mmol/L), mean (± SD) | 4.86 ± 0.24 | 5.06 ± 0.26 | 0.066 |
| α-AMY (U/L), mean (± SD) | 648.3 ± 11.90 | 654.7 ± 13.16 | 0.266 |
| Duct size (mm), mean (± SD) | 2.24 ± 0.16 | 2.16 ± 0.08 | 0.164 |
| Soft gland texture, number (%) | 18 (100) | 18 (100) | — |
| Operative time (min), median (IQR) | 31.5 (28.75-33) | 29 (28-31.25) | 0.138 |

FBG, Fasting blood glucose

**Figure 3.**
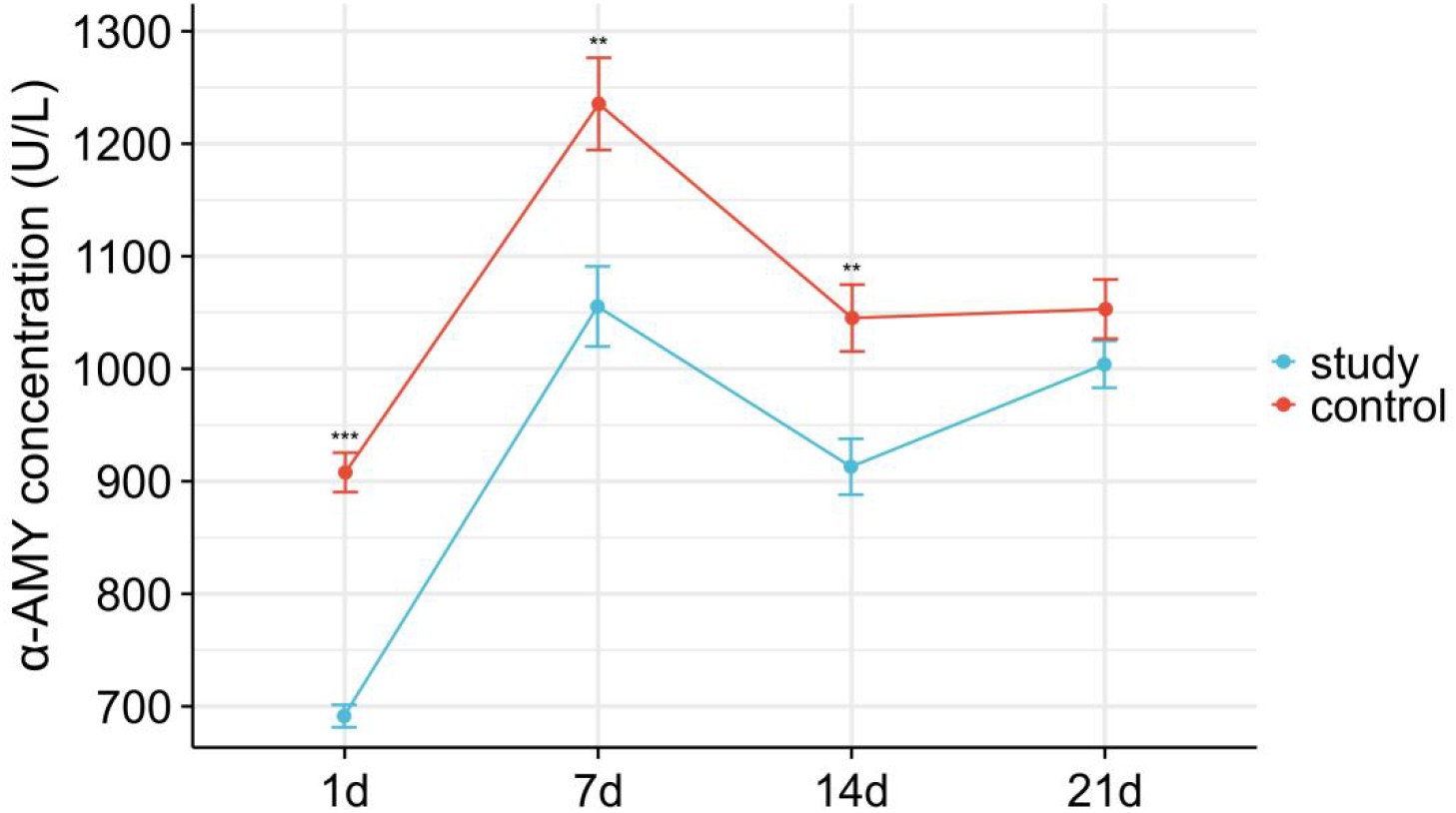
The expression of serum amylase in the two groups of Beagles on the 1st, 7th, 14th, and 21st days after pancreaticointestinal anastomosis.(\*\**p* < 0.01; \*\*\**p* < 0.001)

### 3.2. Postoperative Mortality and Morbidity Rates

The postoperative mortality, morbidity, and causes of death of the dogs in this study were investigated. The surgeries in both the study group and the control group were successfully performed. Two dogs (2/18) in the study group died before the specified sacrifice time, and of these, one died from pulmonary infection, while the other died from gastric perforation. Three dogs (3/18) in the control group died before the specified sacrifice time. The causes of death were anastomotic leakage in 1 case, postoperative bleeding in another case, and pulmonary infection in the third case. The complication rates of the study group and the control group were 5/18 and 7/18, respectively (*p > 0*.*05, χ*^2^ test) (Table 2).

**Table 2.** Comparison of postoperative complications between the two experimental groups of Beagle dogs.

|  | Study group (n=18) | Control group (n=18) | *p* |
| --- | --- | --- | --- |
| Postoperative complications, number (%) | 5 (27.78) | 7 (38.89) | 0.724 |
| Anastomotic leakage | 0 (0.00) | 1 (5.56) |  |
| Pancreatitis | 2 (11.11) | 3 (16.67) |  |
| Pulmonary infection | 1 (5.56) | 1 (5.56) |  |
| Intestinal infection | 1 (5.56) | 1 (5.56) |  |
| Gastric perforation | 1 (5.56) | 0 (0.00) |  |
| Haemorrhage | 0 (0.00) | 1 (5.56) |  |
| Postoperative death, number (%) | 2 (11.11) | 3 (16.67) | >0.999 |

### 3.3. ELISA Detection of Serum TGF-β1, IL-10, TNF-α, and IL-6

The levels of TGF-β1, IL-10, TNF-α, and IL-6 in the serum of dogs in different groups were detected using ELISA kits. The results revealed that the levels of TGF-β1, IL-10, TNF-α, and IL-6 in both groups were increased at all time points. Compared with those in the control group, the increases at some time points in the study group were significant (*\*p < 0*.*05; **p < 0*.*01*) (Figure 4).

**Figure 4.**
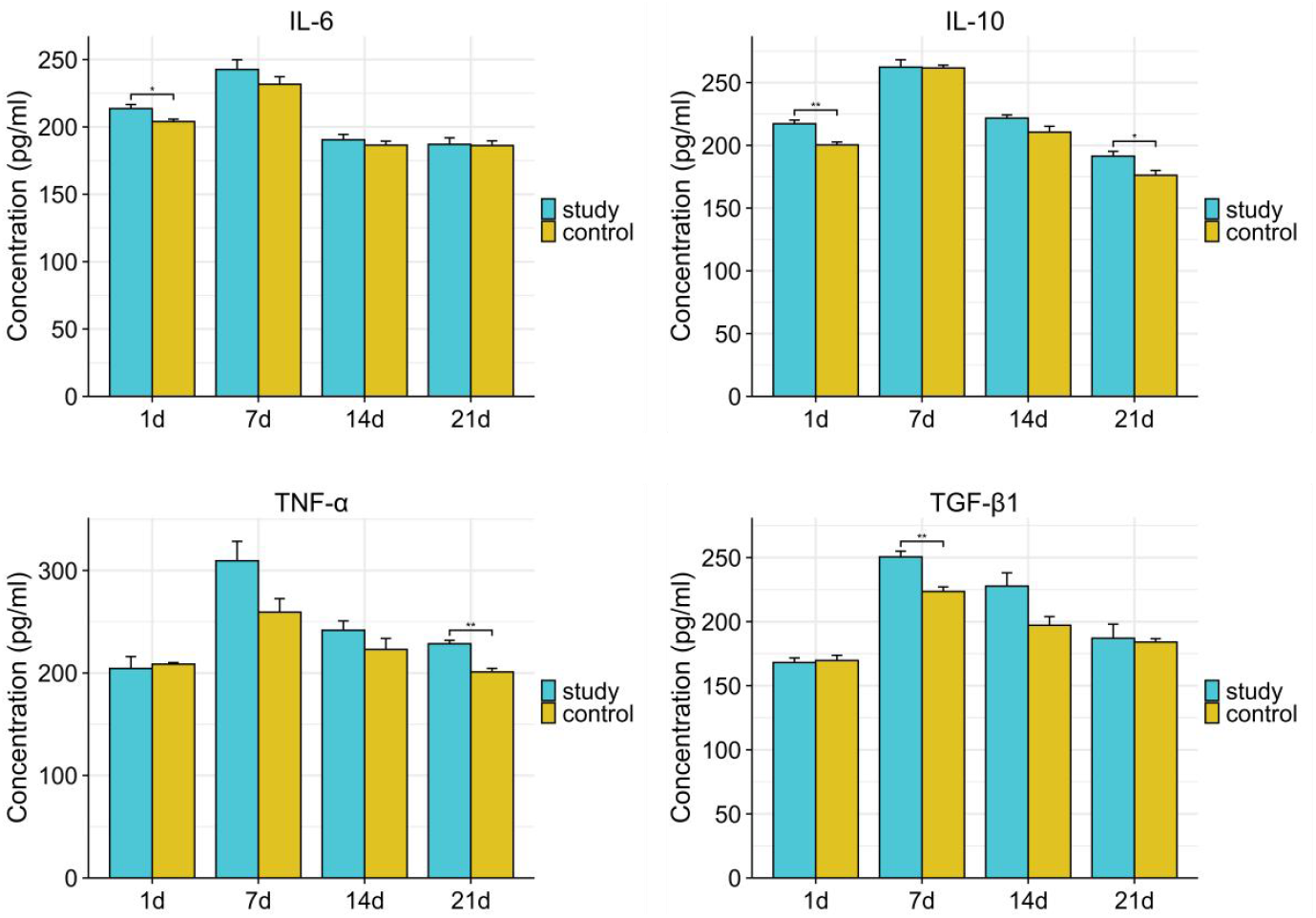
The expression levels of TGF-β1, IL-10, IL-6, and TNF-α in the serum of dogs in the two groups on the 1st, 7th, 14th, and 21st days after pancreaticointestinal anastomosis.(\**p* < 0.05; \*\**p* < 0.01)

### 3.4 Pathological Observation of the Anastomoses

In dogs in the study group, the pancreatic tissue in front of the anastomosis was closely adhered to the jejunal surface at the suture site on the 7th day after surgery. Obvious tissue inflammatory reactions and oedema were also observed. The anastomosis was not completely healed, and a small volume of exudate and a large amount of a pale-yellow necrotic substance could be seen (Figure 5A, A1). In the control group, the suture site of the anastomosis was slightly everted on the 7th day after surgery. A small amount of necrotic tissue was observed on the pancreatic section side, and oedema was obvious (Figure 5B). The pancreatic ducts in both groups were patent and slightly dilated, and both contained clear pancreatic juice. By the 14th day after surgery, the oedema in both groups had significantly subsided. The suture site of the posterior wall of the anastomosis in the study group became thicker, the necrotic tissue at the anastomosis was sloughed off (Figure 5C), and the anastomosis healed well. H&E staining and microscopic examination revealed that the intestinal mucosa and pancreatic duct mucosa at the suture site of the anastomosis were closely connected. A small amount of scar connective tissue was observed under the mucosa (Figure 5C1). Moreover, the tissue at the suture site in the control group was thickened (Figure 5D). Microscopy analysis of H&E-stained sections revealed that the intestinal mucosal epithelial tissue covered the proliferative connective tissue (Figure 5D1). In the study group, the pancreatic cross-section of the anastomosis was completely covered by the intestinal mucosa by the 21st day after surgery. The anterior wall of the pancreaticojejunostomy anastomosis was everted, and the posterior wall protruded in a semicircular shape (Figure 5E). The anastomosis was obviously scarred, and the pancreatic ducts were dilated to different degrees. The intestinal mucosal cells crawled from the periphery to the centre over an increasingly wide range, where they proliferated and differentiated into the lumen. The degree of cell differentiation was high near the anastomosis and approached that of normal intestinal mucosa. On the 21st day after surgery, in the control group, the pancreatic duct was obviously dilated (Figure 5F). Compared with that in the study group, scar hyperplasia at the site of the anastomosis was more obvious. The degree of pancreatic inflammation was severe in 2 dogs. Partial mucosal necrosis and a high level of inflammatory cell infiltration was observed by microscopy.

**Figure 5.**
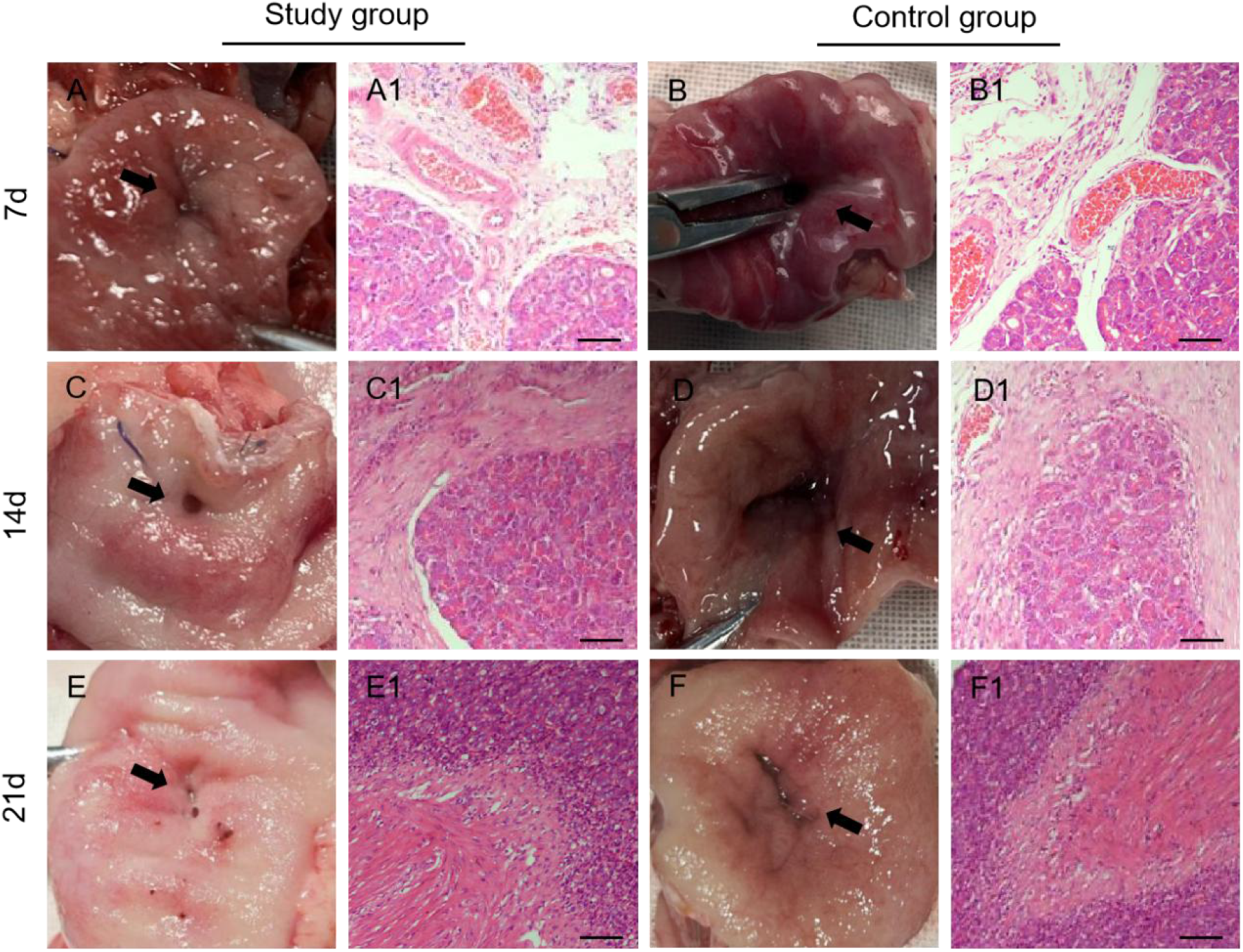
On the 7th day (A, B), 14th day (C, D), and 21st day (E, F) after surgery in the two groups of dogs, the anastomotic site was observed, and H&E staining was performed on the corresponding parts. A-F shows the direct cross-sectional view of the pancreatic intestinal anastomosis site. A1-F1 shows the H&E staining of the corresponding anastomotic site. Arrow heads: Location of pancreatic intestinal anastomosis. Scale bar 200 µm; haematoxylin and eosin.

The immunohistochemical results revealed that at different time points, the protein expression levels of collagen I/III and α-SMA in the anastomosis were increased in both groups. The collagen I protein level gradually increased over time, whereas the collagen III and α-SMA protein levels gradually decreased over time. Compared with those in the control group, the changes in the protein expression levels of collagen I/III and α-SMA in the study group were significant at some time points (*\*p < 0*.*05; ***p < 0*.*001*) (Figure 6). The immunofluorescence results of the pancreaticojejunostomy anastomosis on the 21st day after surgery revealed no significant difference in the fluorescence intensity of the collagen I/III and α-SMA proteins between the study group and the control group (*p > 0*.*05*) (Figure 7).

**Figure 6.**
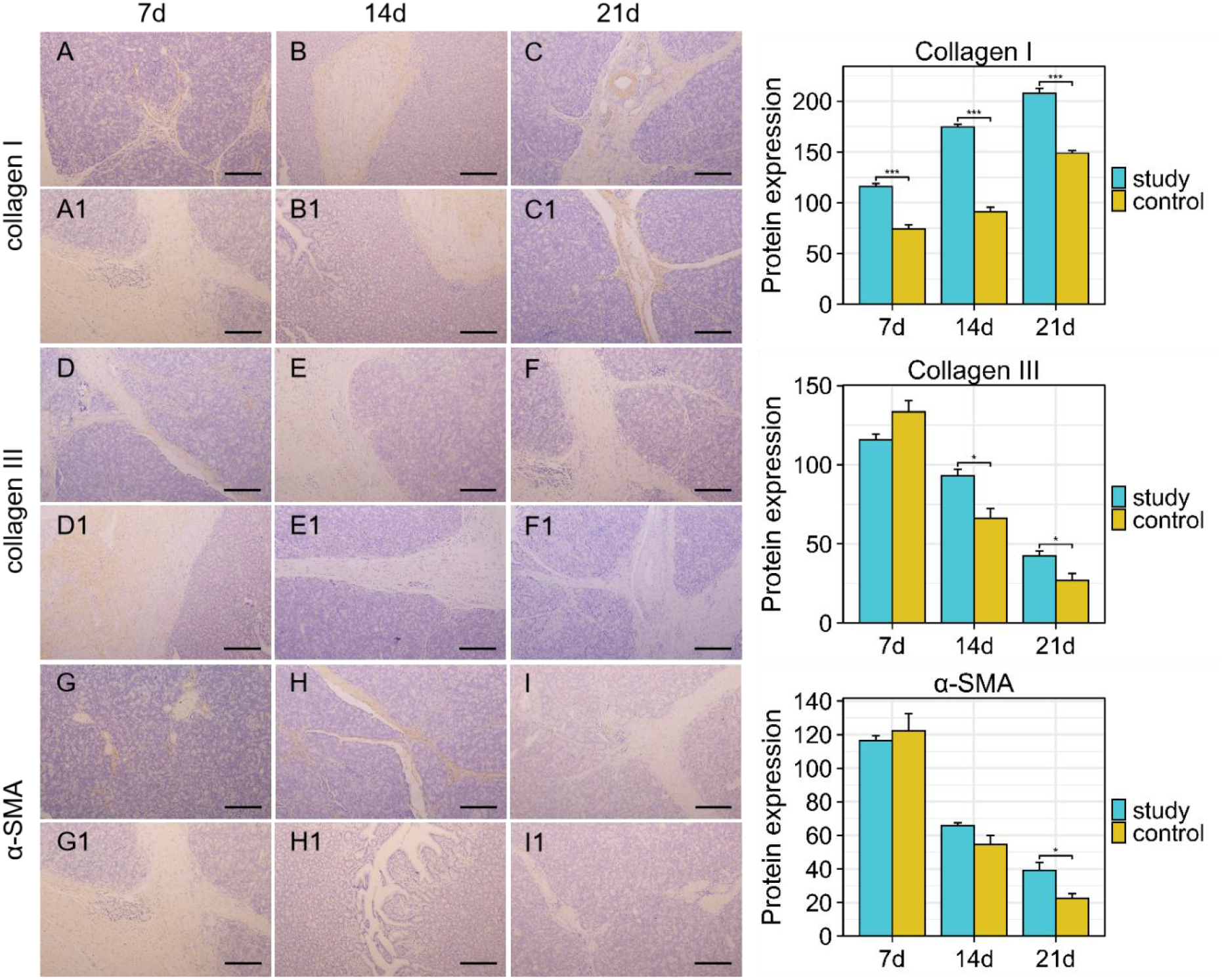
Immunohistochemical staining of the anastomotic site was performed on the 7th, 14th, and 21st days after surgery in the two groups of dogs. The figure shows the protein expression intensity of α - SMA, collagen I, and collagen III at the corresponding sites of the two sets of anastomoses. A-I shows the immunohistochemical staining of the anastomosis in the study group at different time points, while A1-I1 shows the immunohistochemical staining of the anastomosis in the control group at different time points. Scale bar 200 µm; immunohistochemical staining.(\**p* < 0.05; \*\*\**p* < 0.001)

**Figure 7.**
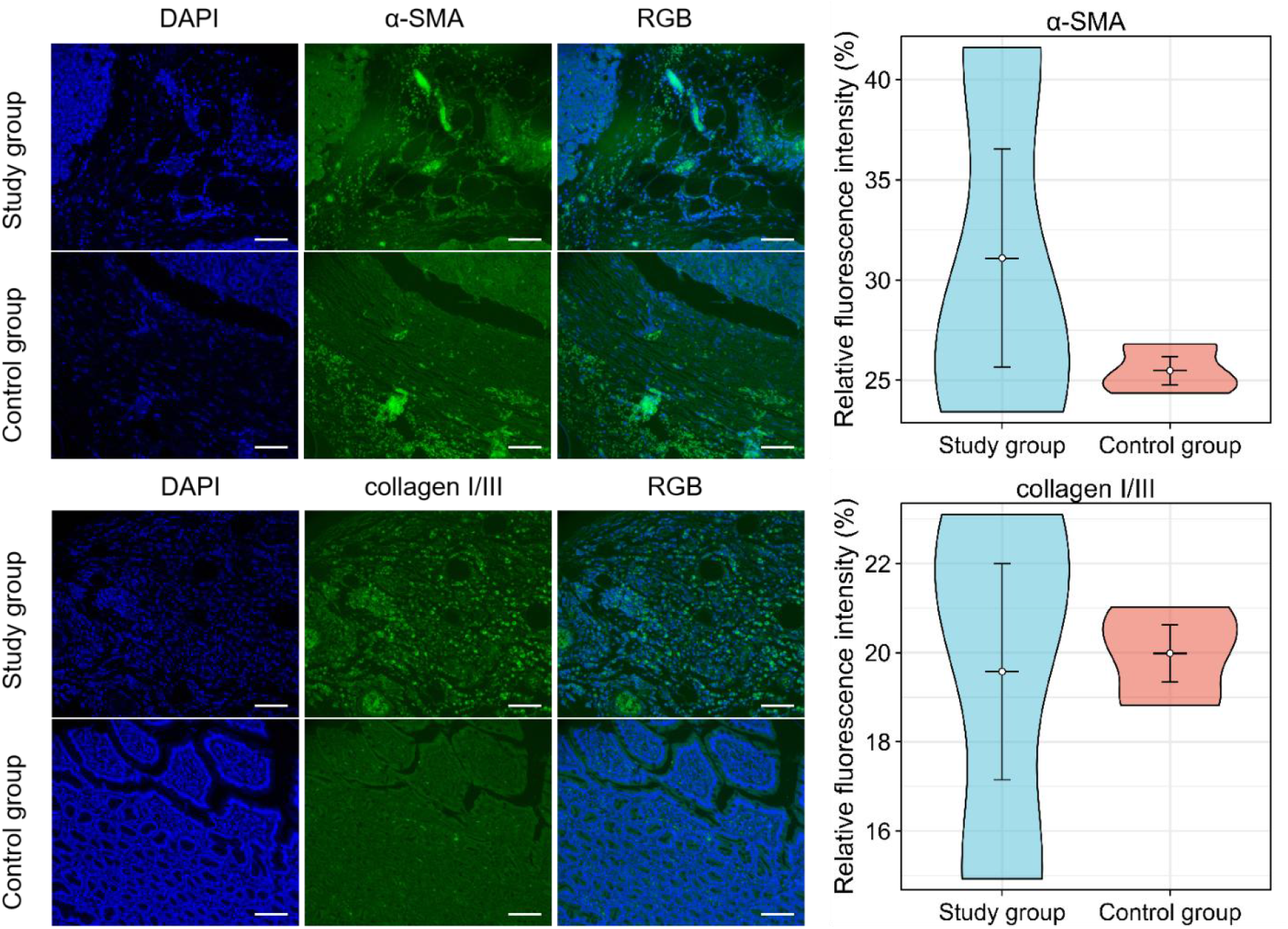
On the 21st day after surgery, sections of the anastomotic site from the two groups of dogs were stained by immunofluorescence. The figure shows the immunofluorescence intensity of α - SMA and collagen I/III at the corresponding sites of the two sets of anastomoses. DAPI: 4’,6-diamidino-2-phenylindole, RGB: In optics, RGB refers to the colour formed by the combination of the three primary colours of red, green, and blue. Scale bar 200 µm; Immunofluorescence staining.

## 4. Discussion

The results of our animal experiments indicate that inflammatory factors play a positive role in the early stage of PJ anastomosis healing by stimulating the secretion and synthesis of collagen. Moreover, for reconstruction after pancreatectomy, pancreaticojejunostomy with mucosal-priority healing has the following advantages: (1) the operation is safe and simple, with relatively low technical difficulty; (2) the anastomosis heals well, and the incidence of anastomotic fistula is low; and (3) the incidence of postoperative bleeding and death is low.

In classic IPJ, the pancreatic stump is completely exposed in the jejunal lumen. Although this procedure avoids the operational difficulties caused by a small pancreatic duct [10], it increases the likelihood of complications resulting from the corrosion of the pancreatic section. The probability of severe complications (grades IIIa, IIIb, IV, and V), including bleeding, that require additional intervention is also significantly greater in IPJ. The smaller the diameter of the pancreatic duct, the greater the degree of surgical difficulty [11]. The PM-PJ does not require a perfect mucosa-to-mucosa anastomosis between the pancreatic duct and the jejunum [12]. This avoids the tearing of small pancreatic ducts caused by close suturing. The connection is sufficient between the pancreatic duct and the jejunum if this area remains unobstructed to allow physiological healing. During anastomosis, the pancreatic tissue on the posterior wall of the pancreatic duct is first sutured to the posterior wall of the jejunum, and then the anterior wall of the jejunum is sutured to the pancreatic tissue on the anterior wall of the pancreatic duct. The surgical field is clear, the steps are simplified, and the procedure is relatively easy.

The animal experiments revealed that in PM-PJ, the posterior wall of the anastomosis essentially healed within 7 days. While one dog in the control group died of anastomotic leakage, none of the dogs in the study group experienced this adverse event. This may be related to the relatively long free pancreas, which affects the blood supply of the stump and leads to poor local blood supply at the anastomosis site [13]. On the 21st day after surgery, the section of the pancreas at the anastomosis in dogs in the study group was covered by proliferative mucosal epithelium, and the unsutured anterior wall of the pancreatic duct had healed in a slightly everted state. This typical healing is only possible when healing of the pancreatic section of the anastomosis is delayed. In the early stage of tissue healing, the wound defect is covered by granulation tissue. In the middle stage, this defect contracts under the action of myofibroblasts. Then, epithelial cells grow towards the centre and finally cover the defect. After epithelialization, the growth of granulation tissue is inhibited, and scar hyperplasia stops [14]. In our study, the early stage of anastomosis healing is the inflammatory reaction stage, during which coagulation, haemostasis, inflammatory exudation and neutrophil infiltration occur and during which inflammatory cells release various inflammatory and tissue growth factors. This process subsequently leads to the tissue-healing stage, in which fibroblasts proliferate, collagen is secreted, capillaries regenerate, and microcirculation is established. The mature stage (remodelling) is the stage at which granulation tissue is remodelled. Due to the unique anatomical structure of the pancreas and intestine and the presence of pancreatic juice, the repair and healing processes after PJ occur according to their own principles and characteristics. When the defect is covered by proliferative mucosal epithelial tissue, the entire anastomosis is epithelialized, thereby preventing further scar hyperplasia. Moreover, the proliferative mucosal epithelium protects the anastomosis tissue from the corrosive effect of pancreatic juice and prevents bleeding caused by tissue corrosion.

Within 21 days of the animal experiment, the serum levels of inflammatory factors, as well as the levels of collagen and α-SMA at the site of the pancreaticojejunostomy anastomosis, increased to varying degrees. At this time point, tissue repair is most easily observable, and the characteristics of skin repair are the most comprehensive. However, the repair processes of all tissues in the body are similar and involve a series of coordinated stages. This process is usually divided into blood clot formation, inflammation, new tissue formation (including epithelial regeneration, granulation tissue formation and contraction), and finally, tissue remodelling and regression. In the early stage, platelet degranulation provides some signals that initiate various aspects of the repair process, including the recruitment of inflammatory cells. The acute inflammatory phase subsides when proinflammatory signals and leukocyte recruitment are reduced, usually through the upregulation of IL-10 and TGF-β [15,16]. Without these factors, the wound cannot heal [17]. In addition, IL-6 is a multifunctional cytokine that can stimulate the secretion of collagen, thus accelerating the repair process [18]. IL-6 promotes the proliferation of fibroblasts and collagen synthesis by activating the JAK/STAT signalling pathway [7]. After pancreaticojejunostomy, plasma components leak into the local anastomosis area and initiate a coagulation reaction to form a blood clot, which provides spatial support for early inflammatory reactions and tissue repair. Moreover, the release of cytokines in the blood clot can regulate the functions of inflammatory cells, fibroblasts, and keratinocytes, among other cells. Subsequently, under stimulation by cytokines such as TNF-α, IL-6, IL-10, and INF-γ, inflammatory cells in the blood vessels migrate away from the local anastomosis area. In the early stage, the cells infiltrating the wound area are mainly neutrophils, which can not only remove local necrotic tissue but also secrete cytokines to induce the migration of other inflammatory cells to the wound[19]. In addition to neutrophils, macrophages also play an immune role in phagocytizing bacteria and necrotic tissue and in antigen presentation. They can also synthesize and release various growth factors, such as PDGF, TGF-β, and VEGF, which play integrative roles in the repair process [20]. Macrophages have an important function in fibroblast signalling through which fibroblasts transform into myofibroblasts and deposit collagen [21].

In this study, although we explored the mechanism by which inflammatory factors affect the migration and repair of intestinal mucosal cells at the PJ anastomosis site, the use of animals in research inevitably has several limitations. According to complication rate statistics, a small sample size may lead to large fluctuations in the incidence rate, and thus it would be impossible to accurately reflect the true complication rates of different surgical methods. This study only observed the effects of inflammatory factors on anastomosis healing within 21 days after surgery, which may not be sufficient to fully capture the entire process of anastomosis healing and its long-term effects. Although inflammatory factors stimulate the secretion and synthesis of collagen in the early stage of PJ anastomosis healing, the signalling pathways through which inflammatory factors specifically act and how they interact with other intracellular molecules to affect the migration and repair of intestinal mucosal cells remain unknown.

## 5. Conclusion

This study revealed that inflammatory factors stimulate the secretion and synthesis of collagen in the early stage of PJ anastomosis healing, which is helpful for anastomosis healing. The PM-PJ procedure is safe and simple and can effectively reduce the degree of technical difficulty. Moreover, this surgical method can ensure good anastomosis healing and significantly reduce the incidence of anastomotic fistula, postoperative bleeding, and death, providing a new approach for reconstruction after pancreatectomy.

## Ethics approval and consent to participate

Ethical approval for this study (number: IACUC-20210303-55) was provided by the Ethical Committee of Xinjiang Medical University.

## Consent for publication

Not applicable.

## Availability of data and materials

The datasets used and/or analysed during the current study are available from the corresponding author on reasonable request.

## Competing interests

The authors declare that they have no competing interests.

## Funding

This work has been funded by National Natural Science Foundation of China (Award Number: 82160497) and Natural Science Foundation of Xinjiang Uygur Autonomous Region (Award Number: 2021D01C424).

## Authors’ contributions

Shixing Wu Tigu A and Bolin Zhang Li wrote the manuscript; Cheng Wang and Jiangang Li performed data aggregation; Junxiang Zhang and Shouchao Zhang collected clinical data; Cheng Geng performed Conceptualization; Xinjian Xu Performed Methodology and obtaining funding.

## Acknowledgements

Not applicable.

## Notes

### Competing Interest Statement

The authors have declared no competing interest.

